# Distinguishing COVID-19 infection and vaccination history by T cell reactivity

**DOI:** 10.1101/2021.12.15.472874

**Authors:** Esther Dawen Yu, Eric Wang, Emily Garrigan, Benjamin Goodwin, Aaron Sutherland, James Chang, Rosa Isela Gálvez, Jose Mateus, Stephen A. Rawlings, Davey M. Smith, April Frazier, Daniela Weiskopf, Jennifer M. Dan, Shane Crotty, Alba Grifoni, Ricardo da Silva Antunes, Alessandro Sette

**Author notes:** These authors contributed equally (Ricardo da Silva Antunes, Alessandro Sette). Correspondence should be addressed to Ricardo da Silva Antunes, La Jolla Institute for Immunology, La Jolla, CA 92037, USA, and Alessandro Sette, La Jolla Institute for Immunology, La Jolla, CA 92037, USA.

## Abstract

SARS-CoV-2 infection and COVID-19 vaccines elicit memory T cell responses. Here, we report the development of two new pools of <u>E</u>xperimentally-defined T cell epitopes derived from the non-spike <u>R</u>emainder of the SARS-CoV-2 proteome (CD4RE and CD8RE). The combination of T cell responses to these new pools and Spike (S) were used to discriminate four groups of subjects with different SARS-CoV-2 infection and COVID-19 vaccine status: non-infected, non-vaccinated (I−V−); infected and non-vaccinated (I+V−); infected and then vaccinated (I+V+); and non-infected and vaccinated (I−V+). The overall classification accuracy based on 30 subjects/group was 89.2% in the original cohort and 88.5% in a validation cohort of 96 subjects. The T cell classification scheme was applicable to different mRNA vaccines, and different lengths of time post-infection/post-vaccination. T cell responses from breakthrough infections (infected vaccinees, V+I+) were also effectively segregated from the responses of vaccinated subjects using the same classification tool system. When all five groups where combined, for a total of 239 different subjects, the classification scheme performance was 86.6%. We anticipate that a T cell-based immunodiagnostic scheme able to classify subjects based on their vaccination and natural infection history will be an important tool for longitudinal monitoring of vaccination and aid in establishing SARS-CoV−2 correlates of protection.

## INTRODUCTION

Immune memory against severe acute respiratory syndrome coronavirus 2 (SARS-CoV−2) is associated with cellular and humoral adaptive immunity (Painter et al., 2021; Rydyznski Moderbacher et al., 2020; Sette and Crotty, 2021). Correlates of protection from infection and symptomatic disease have not been firmly established (Feng et al., 2021; Koup et al., 2021; Krammer, 2021a, b), and may require comprehensive assessment of antibody titers and levels of effector and memory B and T cell responses. Broad measurement of T cell responses is hindered by the lack of immunodiagnostics tools with effective predictive power able to discriminate pre-existing immunity, vaccination, and infection (Ogbe et al., 2021; Peeling and Olliaro, 2021; Sekine et al., 2020; Vandenberg et al., 2021).

While SARS-CoV−2 T cell responses are detected in nearly all COVID-19 convalescent individuals (Grifoni et al., 2020b; Le Bert et al., 2020; Tarke et al., 2021a), they are also found in 20-50% of unexposed individuals (Mateus et al., 2020; Sette and Crotty, 2020; Tarke et al., 2021a). However, recent evidence suggests that SARS-CoV−2 infection generates a largely novel repertoire of T cells, with over 80% of the epitopes not recognized in unexposed donors (Mateus et al., 2020; Tarke et al., 2021a). In addition, mRNA or viral vector vaccines boost the spike protein-specific immune responses in both unexposed and convalescent individuals without affecting the responses to non-spike SARS-CoV−2 components (Bertoletti et al., 2021; Lozano-Ojalvo et al., 2021; Mateus et al., 2021). Further complexity is associated with evaluating responses in subjects previously infected and subsequently vaccinated, and conversely, previously vaccinated and subsequently infected (breakthrough infection) (Goel et al., 2021; Lucas et al., 2021; Niessl et al., 2021; Rovida et al., 2021). It is therefore a possibility to develop epitope pools based on spike and the rest of the genome reactivity as a tool to discriminate subjects based on their vaccination and natural infection history.

We have shown that SARS-CoV−2 specific T cells can be detected and quantitated using peptide pools in various T cell assays (da Silva Antunes et al., 2021; Dan et al., 2021; Grifoni et al., 2020a; Mateus et al., 2020; Tarke et al., 2021b) which have proven useful to derive information about the kinetics and magnitude of SARS-CoV−2 specific T cell responses in both COVID-19 infection and vaccination (Dan et al., 2021; Mateus et al., 2021). Subsequent studies detailed the repertoire of epitope specificities recognized in a cohort of COVID-19 convalescent subjects (Tarke et al., 2021a). More recently, a meta-analysis of experimental curated data from the Immune Epitope Database (IEDB) revealed a large repertoire of over 1400 epitopes defined in 25 different studies (Grifoni et al., 2021). Here, we used this information to develop SARS-CoV−2-specific peptide pools optimized for broader epitope repertoire and wider HLA coverage for both CD4+ and CD8+ T cell responses. Accordingly, two pools of <u>E</u>xperimentally defined epitopes derived from the non-spike <u>R</u>emainder of the SARS-CoV−2 proteome, (CD4RE and CD8RE) were established.

Several platforms and strategies have been developed to assess T cell responses in both vaccinated or infected individuals, using different read-outs and technologies, such as cytokine release assays (ELISPOT, or ELISA) (Krishna, Preprint; Kruse et al., 2021; Martínez-Gallo, 2021; Murugesan, Preprint; Tan et al., 2021; Tormo, Pre-proof) or flow cytometry-based assays (Blast transformation, or intracellular cytokine staining (ICS)) (Lind Enoksson et al., 2021; Zelba et al., 2021). These assays mainly rely on the characterization of responses to the spike or nucleocapsid antigens, and therefore do not address the entire SARS-CoV−2 proteome and the remarkable breadth of T cell responses against this pathogen (Grifoni et al., 2021).

In this study, we developed an immunodiagnostic T cell assay using a pool of overlapping peptides spanning the entire spike protein in combination with experimentally defined non-spike pools to classify subjects based on their vaccination and natural infection history. This tool showed a highly predictive power to discriminate responses based on distinctive COVID-19 immune profiles, including breakthrough infections, and clinical applicability demonstrated by using a validation cohort, different vaccine platforms, and assessment of responses at different lengths of time post-infection/post-vaccination.

## RESULTS

### Cohorts associated with known infection and vaccination history

239 participants were enrolled in the study and classified into five groups based on known vaccination and clinical history: (50 non-infected, non-vaccinated (I−V−); 50 infected and non-vaccinated (I+V−); 66 infected and then vaccinated (I+V+); 50 non-infected and vaccinated (I−V+); and 23 vaccinated and then infected (V+I+). An overview of the characteristics from all the participants is provided in **Table 1**. For the I+V−, I+V+ and V+I+ groups, SARS-CoV−2 infection was determined by PCR test during the acute phase of infection or verified by serological detection of antibodies against the SARS-CoV−2 Spike protein RBD region at the time of blood donation.

**Table 1.** Description of donor cohort characteristics and demographics.

| Cohort Name | I-V- | I+V- | I-V+ | I+V+ | V+I+ |
| --- | --- | --- | --- | --- | --- |
| **Number of donors** | 50 | 50 | 66 | 50 | 23 |
| **Gender (M/F)** | (26, 24) | (21, 29) | (28, 38) | (23, 27) | (7, 16) |
| **Median age (years)** | 25 (17-64) | 42 (19-67) | 40 (21-74) | 38 (21-73) | 30 (22-68) |
| **Race** |  |  |  |  |  |
| White (n (%)) | 32 (64%) | 37 (74%) | 38 (58%) | 39 (78%) | 15 (65%) |
| Hispanic/Latino (n (%)) | 8 (16%) | 7 (14%) | 11 (17%) | 6 (12%) | 5 (22%) |
| Asian (n (%)) | 8 (16%) | 4 (8%) | 16 (24%) | 3 (6%) | 3 (13%) |
| Black (n (%)) | 2 (4%) | 2 (4%) | 2 (3%) | 2 (4%) | 0 (0%) |
| **Sample collection date** | 2013-2019 | 2020-2021 | 2021 | 2021 | 2021 |
| **COVID-19 vaccination status** | none | none | Vaccinated | Vaccinated | Vaccinated |
| Pfizer (n (%)) | - | - | 30 (45%) | 25 (50%) | 15 (65%) |
| Moderna (n (%)) | - | - | 36 (55%) | 25 (50%) | 8 (35%) |
| Days from 2<sup>nd</sup> dose of vaccination | - | - | 16 (13-190) | 32 (7-188) | 163(55-271) |
| **SARS-CoV-2 status** | Ab(-) | Ab(+) or PCR(+) | Ab(+) and PCR(-) | Ab(+) or PCR(+) | PCR(+) |
| **SARS-CoV-2 PCR (n (%))** |  |  |  |  |  |
| Positive | 0 (0) | 47 (94%) | 0 (0) | 45 (90%) | 23 (100%) |
| Unknown | - | 3 (6%) | - | 5 (10%) | - |
| **Spike (S) antibody response (n (%))** |  |  |  |  |  |
| Median | 3.0 | 191.8 | 4157.0 | 4654.0 | 8783.0 |
| Range | 3.0-23.9 | 3.0-7326.0 | 410.6-32033.0 | 159.2-25876.0 | 2165.0-35319.0 |
| **Nucleocapsid (N) antibody response (n (%))** |  |  |  |  |  |
| Median | 4.9 | 177.9 | 16.3 | 73.2 | 241.5 |
| Range | 3.0-339.8 | 3.0-11755.0 | 3.0-109.0 | 3.0-1873.0 | 72.6-5044.0 |
| **Post symptom onset (days)** |  |  |  |  |  |
| Median | - | 119 | - | 354 | 32 |
| Range | - | 20-308 | - | 57-508 | 18-93 |
| **Symptoms (n (%))** |  |  |  |  |  |
| Asymptomatic | - | 0 (0%) | - | 0 (0%) | 0 (0%) |
| Mild | - | 44 (88%) | - | 45 (90%) | 23 (100%) |
| Moderate | - | 3 (6%) | - | 3 (6%) | 0 (0%) |
| Severe | - | 3 (6%) | - | 2 (4%) | 0 (0%) |

Summary of donor characteristics.
Non-infected, non-vaccinated (I-V-); infected and non-vaccinated (I+V-); infected and then vaccinated (I+V+); non-infected and vaccinated (I-V+); and vaccinated and then infected (V+I+)

The study primarily consisted of subjects recruited in San Diego, California (see material and methods for more details). Among individuals with history of COVID-19 disease, the majority were symptomatic mild disease cases, owing to the nature of the study recruitment design. Specifically, 44 donors (88%) for I+V−, 45 donors (90%) for I+V+, and 23 donors (100%) for V+I+ had mild symptoms, 3 donors (6%) of I+V− and I+V+ groups had moderate symptoms, and 3 (6%) and 2 donors (4%) from the I+V− and I+V+ groups, respectively, had severe symptoms. The median days of blood collection post symptom onset (PSO) were 119 (20-308), 354 (57-508) and 32 (18-93) for I+V−, I+V+ and V+I+ groups respectively. For the I−V+, I+V+ and V+I+ groups, the vaccinated subjects received two doses of mRNA vaccines BNT162b2 (Pfizer/BioNTech) or mRNA-1273 (Moderna), as verified by vaccination records and positive plasma SARS-CoV−2 spike protein RBD IgG titers. Similar distribution of Pfizer or Moderna administered vaccines (45%-55%) were present in vaccinated subjects from either the I−V+ or I+V+ group, while in the V+I+ group, 15 (65%) subjects had received the BNT162b2 vaccine, and 8 (35%) the mRNA-1273 vaccine.

The median days of blood collection post second dose of vaccination (PVD) were 16 (13-190), 32 (7-188) and 163 (55-271) for I−V+, I+V+ and V+I+ groups, respectively. All the I−V−subjects were collected before the attributed pandemic period (2013-2019) and confirmed seronegative with undetectable SARS-CoV−2 Spike protein RBD IgG titers. In all cohorts, the median ages were relatively young (25 (17-64), 42 (19-67), 40 (21-74), 38 (21-73), 30 (22-68) for I−V−, I+V−, I−V+, I+V+ and V+I+ groups respectively), with the female gender well represented and different ethnicities represented. In our study, participants were further divided in an exploratory cohort (120 donors, **Table S1**), an independent validation cohort (96 donors, **Table S2**) and a third cohort of breakthrough infections (V+I+; 23 donors, **Table 1**).

### Differential SARS-CoV−2 CD4+ T cell responses in unexposed, convalescent, and vaccinated subjects

To detect SARS-CoV−2 T-cell reactivity, we previously routinely utilized a pool of overlapping peptides spanning the entire spike (S) sequence (253 peptides) and a pool of predicted HLA Class II binders from the <u>R</u>emainder (R) of the genome (CD4R; (221 peptides) (Grifoni et al., 2020b) (**Table S3)**. Here to further optimize detection of non-Spike reactivity, we designed epitope pools based on <u>E</u>xperimentally (E) defined epitopes, from the non-spike sequences of the SARS-CoV−2 proteome. The CD4RE and CD8RE megapools (MP) consisted of 284 and 621 peptides respectively (**Table S3 and S4**). A pool of epitopes derived from an unrelated ubiquitous pathogen (EBV) (Carrasco Pro et al., 2015) was used as a specificity control (**Table S3)**.

T cell reactivity was assessed by the Activation Induced Marker (AIM) assays (da Silva Antunes et al., 2021) and data represented as either absolute magnitude or stimulation index (SI). As shown in **Figure 1A** SARS-CoV−2-specific CD4+ T cell responses were detected in all convalescent and/or vaccinated individuals and approximately 50% of non-infected, non-vaccinated individuals. Similar results were observed when responses were plotted as SI (**Figure 1B**). Unexposed subjects were associated with significantly lower reactivity as compared to all the other groups (*p*-values ranging 1.3e-7 to 1.0e-15) and convalescent and vaccinated (I+V+) subjects exhibited higher responses than convalescent (I+V−) subjects (p=0.02 and p=0.04 for absolute magnitude and SI, respectively) or vaccinated (I−V+) subjects (p=0.01 and p=0.02 for absolute magnitude and SI, respectively) (**Figure 1A,B**). Importantly, CD4RE responses were able to differentiate convalescent subjects (I+V− or I+V+) from unexposed and vaccinated (I−V+) subjects with *p*-values ranging 5.6e-8 to 5.7e-12 and vaccinated (I−V+) from infected and vaccinated (I+V+) subjects (p=1.4e-11 and p=1.1e-11 for absolute magnitude and SI, respectively) (**Figure 1A,B**). No statistically significant difference in EBV reactivity was observed when the four groups were compared, as expected (**Figure 1A,B)**.

**Figure 1.**
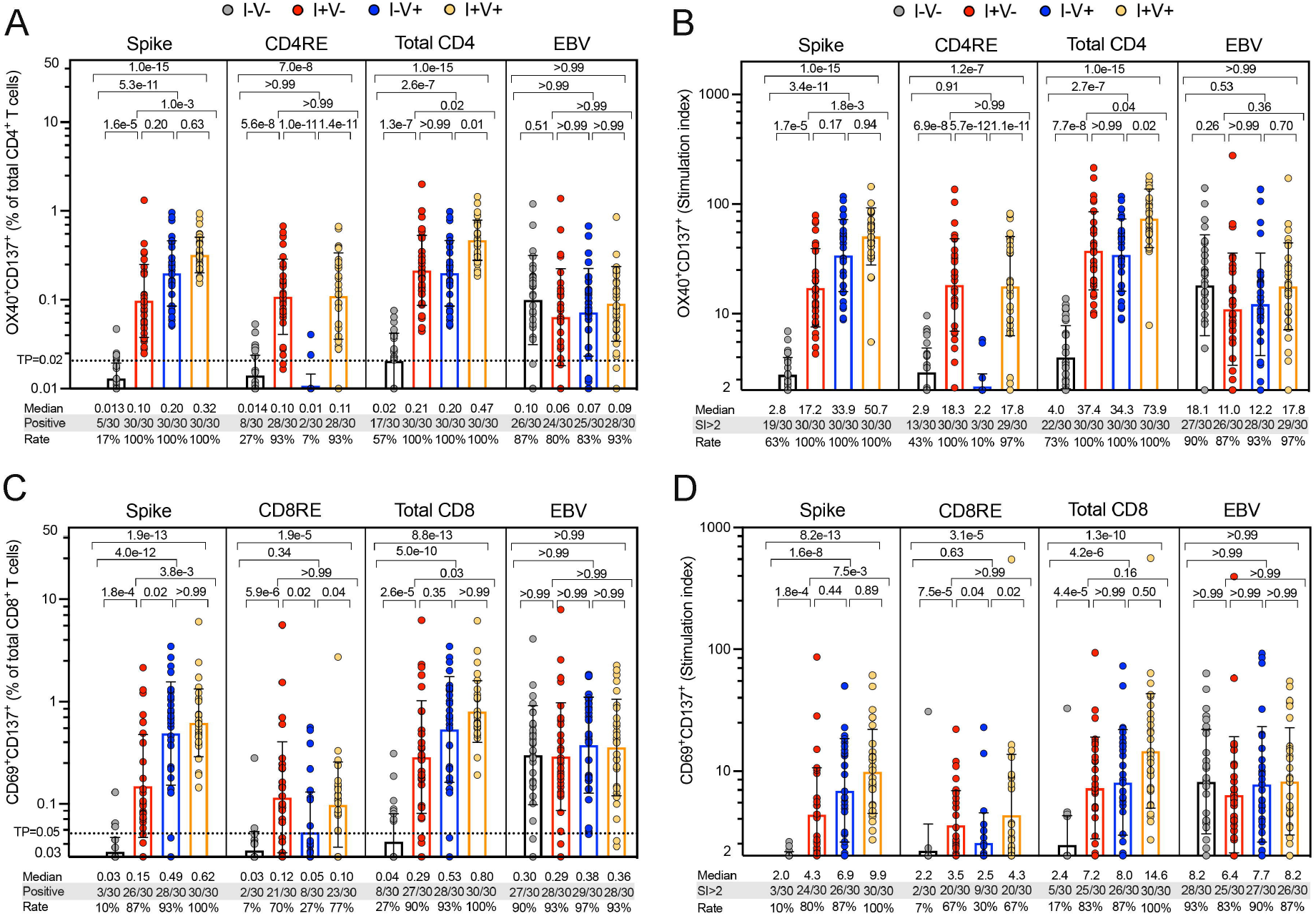
SARS-CoV−2-specific CD4+ and CD8+ T cell responses in the study groups. SARS-CoV−2-specific T cell responses were measured as percentage of AIM+ (OX40+CD137+) CD4+ T cells or AIM+ (CD69+CD137+) CD8+ T cells after stimulation of PBMCs with peptides pools encompassing spike only (Spike) MP or the experimentally defined CD4RE and CD8RE MPs representing all the proteome without spike. EVB MP was used as a control. Graphs show individual response of spike, CD4RE or CD8RE and the combination of both (Total CD4+ or Total CD8+) plotted as background subtracted (**A, C**) or as SI (**B, D**) against DMSO negative control. Geometric mean for the 4 different groups is shown. Kruskal-Wallis test adjusted with Dunn’s test for multiple comparisons was performed and *p* values < 0.05 considered statistically significant. I−V−, unexposed and unvaccinated (n=30); I+V−, infected and non-vaccinated (n=30); I+V+, infected and then vaccinated (n=30); I−V+, non-infected and vaccinated (n=30). Threshold of positivity (TP) is indicated. Median response, and the number or percentage of positive responding donors for each group is shown.

### Differential SARS-CoV−2 CD8+ T cell and IFNγ FluoroSpot responses in unexposed, convalescent, and vaccinated subjects

SARS-CoV−2 specific CD8+ T cell responses were also broadly detected among all the cohorts studied. CD8+ T cell responses were detected in 90-100% of the convalescent and/or vaccinated individuals and approximately in 1/4 of non-infected, non-vaccinated individuals (**Figure 1C)**. Similar responses were observed when plotted as SI (**Figure 1D**). As observed for CD4+ T cell responses, unexposed subjects (I−V−) were discriminated from all the other groups (*p*-values ranging 2.6e-5 to 8.8e-13) and I+V+ infected/vaccinated subjects exhibited higher responses than I+V−convalescent (p=0.03 and p=0.16 for absolute magnitude and SI respectively). Identical results were observed parsing spike-specific responses with CD8RE able to differentiate convalescent (I+V−) from unexposed and vaccinated (I−V+) subjects (*p*-values ranging 0.02 to 5.9e-6) and vaccinated from infected/vaccinated (I+V+) subjects (p=0.04 and p=0.02 for absolute magnitude and SI, respectively) (**Figure 1C,D**). No statistically significant difference in EBV reactivity was observed when the four groups were compared (**Figure 1C,D)**.

In parallel, an IFNγ FluoroSpot assay was also employed to evaluate the CD4+ and CD8+ T cell responses using a threshold of 20 IFNγ spot forming cells (SFC) per million PBMC. Responses were detected in many infected or vaccinated individuals and similar results were observed for Spike, CD4RE or CD8RE when considering both the absolute magnitude or stimulation index, albeit as expected, with lower sensitivity and specificity compared to AIM (**Figure S1)**.

### Improved performance of the CD4RE pool based on experimentally defined epitopes

Results from both AIM and IFNγ FluoroSpot assay demonstrated that the newly developed CD4RE pool had both improved sensitivity and specificity, compared to the previously used CD4R pool of predicted epitopes (**Figure S2A**). In more detail, higher positive CD4+ T cell responses in I+V− (28/30 (93%) vs 26/30 (87%), *p* = 2.0e-4) and I+V+ (28/30 (93%) vs 23/30 (77%), *p* = 5.0e-6), and lower non-specific response in I−V− (8/30 (27%) vs 14/30 (47%), *p* = 0.037) and I−V+ (2/30 (7%) vs 4/30 (13%), *p* = 0.031) were detected using CD4RE when compared to CD4R in the AIM assay (**Figure S2A**). Similar results were shown by IFNγ FluoroSpot, assay albeit with lower sensitivity compared to AIM (**Figure S2B**). These results demonstrate that the use of experimentally defined, as opposed to predicted epitopes provides higher signal in SARS-CoV−2 exposed subjects, while lowering responses from non-exposed subjects.

### Classification of subjects with different exposure history based on Spike and CD4RE reactivity

We reasoned that unexposed (I−V−) subjects would be unreactive to experimentally defined SARS-CoV−2 peptide pools, while uninfected vaccinated (I−V+) subjects should react only to the S pool. We further reasoned that infected (I+V−) subjects should recognize both S and CD4RE, but infected and vaccinated (I+V+) subjects would have a higher relative S reactivity than infected only (I+V−), as is often the case with hybrid immunity (Crotty, 2021), due to exposure to S twice, once during infection and the other during vaccination.

As shown in **Figure 2A**, spike- and CD4RE-specific CD4+ T cell responses derived from the AIM assay were arranged in a two-dimensional plot. Each dot represents a single subject from a total of 120 donors (30 for each of the 4 groups, **Table S1**). Optimal cutoffs were established to discriminate the four groups and the positive predictive value (PPV), negative predictive value (NPV), sensitivity and specificity were calculated for each individual group.

**Figure 2.**
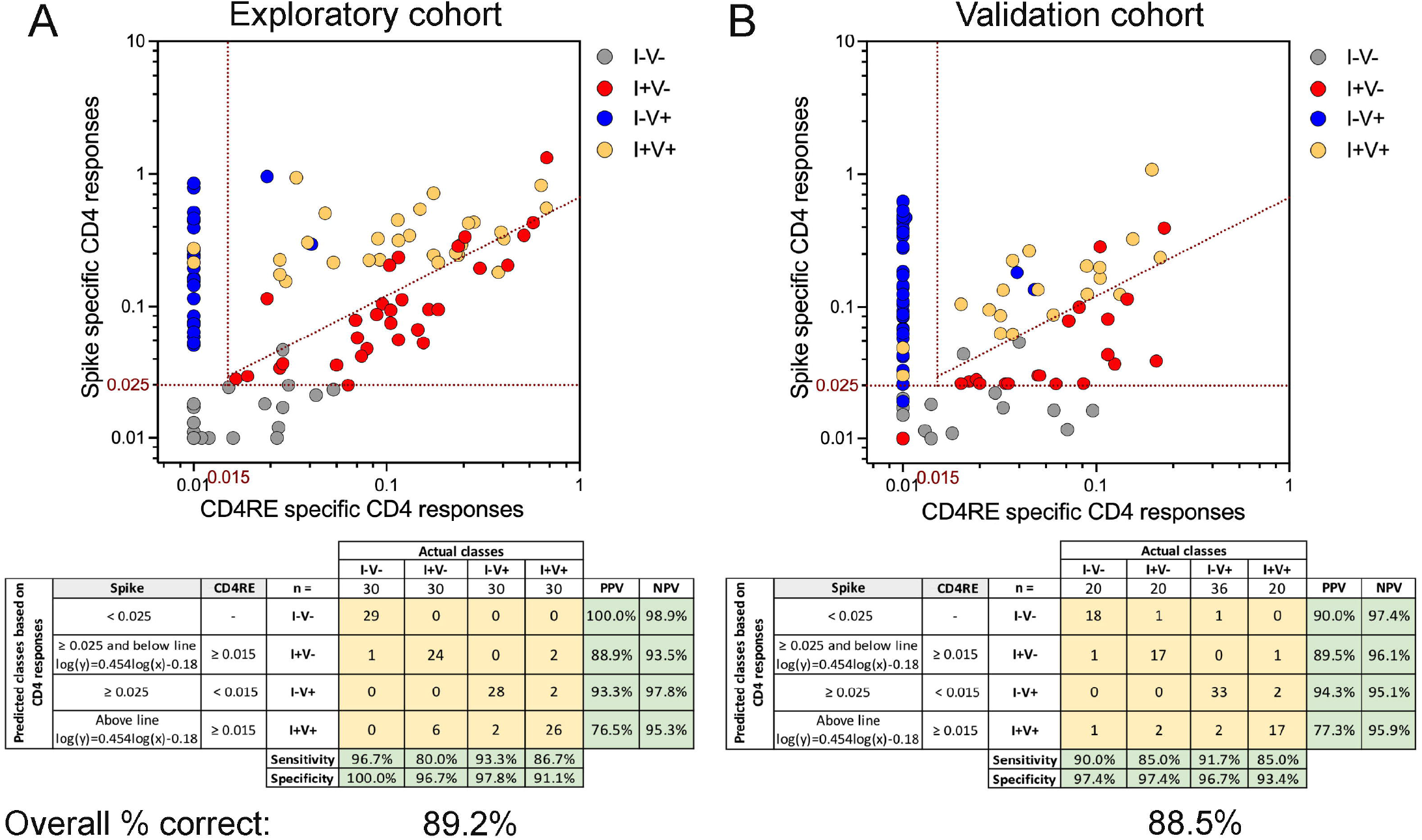
COVID-19 clinical classification scheme using SARS-CoV−2-specific CD4+ T cell responses. CD4+ T cell responses to spike and CD4RE MPs were measured as percentage of AIM+ (OX40+CD137+) CD4+ T cells and plotted in two dimensions as absolute magnitude in order to discriminate the 4 study groups with known COVID-19 status of infection, and/or vaccination in 2 independent cohorts: (**A**) Exploratory cohort (n=120) and (**B**) Validation cohort (n=96). I−V−, unexposed and unvaccinated (n=30 and n=20); I+V−, infected and non-vaccinated (n=30 and n=20); I+V+, infected and then vaccinated (n=30 and n=20); I−V+, non-infected and vaccinated (n=30 and n=36). Red dotted lines indicate specific cutoffs. Table inserts depict the diagnostic exam results in 4×4 matrix. Sensitivity, specificity, PPV, NPV and overall percentage of subjects classified correctly is shown.

Subjects with spike responses lower than 0.025% were classified predictively as unexposed (I−V−) (**Figure 2A**). 29 out of 29 subjects with responses matching this criterion were correctly classified (100% of PPV), while nearly all the actual I−V−subjects (29 out of 30) were found to be associated with responses below the threshold, corresponding to a sensitivity of 96.7 % (**Figure 2A, grey circles)**. Subjects with spike responses greater than 0.025% and CD4RE responses lower than 0.015% were classified predictively as I−V+. Twenty-eight out of 30 subjects with responses matching this threshold were correctly classified (93.3% of PPV), and 28 out of the 30 I−V+ subjects detected within this threshold (93.3% of sensitivity) (**Figure 2A, blue circles)**.

Lastly, subjects with spike and CD4RE responses above 0.025% and 0.015% respectively, and above or below a diagonal line (log(y)=0.454log(x)-0.18) were classified as I+V+ or I+V−respectively. 24 out of 27 subjects with responses matching the lower compartment (I+V−) were correctly classified (88.9% of PPV) while 24 out of the 30 I+V−subjects were found to be associated with this threshold (80 % of sensitivity) (**Figure 2A, red circles)**. Conversely, the majority of subjects (26 out of 34) with responses matching the upper compartment (I+V+) were correctly classified (76.5% of PPV), while 26 out of the 30 I+V+ subjects studied were found to be associated with this threshold, corresponding to a sensitivity of 86.7 % (**Figure 2A, yellow circles)**. Further statistical examinations to assess the robustness of the classification scheme as a potential diagnostic test were performed, specifically assessments of specificity and negative predictive value (PPV). High specificity and NPV were observed for each individual group with a range of 91.1-100% and 93.5-98.9% respectively (**Figure 2A**). In summary, good PPV, NPV, sensitivity and specificity values were observed across all the groups with an overall classification accuracy of 89.2%.

### Validation of the classifier in an independent cohort

To confirm the accuracy of this classification scheme, we assessed CD4+ T cell responses in an independent validation cohort of 96 donors (20 for I−V−, I+V−, I+V+, and 36 for I−V+; **Table S2**). As shown in **Figure 2B**, using the same cutoffs as described above for spike and CD4RE responses, similar PPV, NPV, sensitivity and specificity to the experimental cohort was observed across all the groups with an overall classification accuracy of 88.5%. To further validate the robustness of this classification scheme, the same data (**Figure 2**) was plotted as a function of the stimulation index (**Figure S3)**. Strikingly, these results paralleled the observations using the absolute magnitude of responses, with a similar overall classification accuracy (86.7% and 85.4% for the exploratory and validation cohorts, respectively).

Applying the same classification scheme using either absolute magnitude or stimulation index for IFNγ responses yielded an overall classification accuracy of 72.5% and 60.0% respectively (**Figure S4)**. A lower accuracy was observed when CD8+ T cell responses from AIM assay were analyzed, as compared to CD4+ T cell responses (data not shown). Overall, these results demonstrate the feasibility of assessing CD4+ T cell responses in an integrated classification scheme as a clinical immunodiagnostic tool. Importantly, it also displays the potential for a diagnostic application to discriminate previous undetected infection in vaccinated individuals.

### The classification scheme is applicable to different vaccine platforms, and different lengths of time post-infection/post-vaccination

To gain further insights on the applicability of the classification scheme, we sought to further test and validate this tool when considering different types of vaccines, and longer timepoints post-symptom onset (PSO) or post-vaccination. First, we looked at the response classification as a function of whether vaccinated subjects received BNT162b2 or mRNA-1273 vaccines. As shown in **Figure 3A** the overall classification accuracy when using the different mRNA vaccines was of 89.7%. Specifically, both vaccines showed similar magnitude for both total CD4+ and CD8+ T cell responses in the I−V+ or I+V+ groups (**Figure S5A** and **B**). The accuracy of the classification scheme for the different types of vaccines in the combined I−V+ or I+V+ groups was almost identical (88.5% and 90.9% for the mRNA-1273 and BNT162b2 vaccines, respectively) (**Table 2**).

**Table 2.** Summary of the % correct and applicability of classification scheme.

| **Variable** | **Group** | **AIM assay<sup>\*</sup>**<br>**% correct** |
| --- | --- | --- |
| Type of vaccine | mRNA-1273 | 88.5 |
|  | BNT162b2 | 90.9 |
| Days PSO | Early | 83.7 |
|  | Late | 84.3 |
| Days post-vaccination | Early | 91.2 |
|  | Late | 87.5 |
| <sup>*</sup> CD4<sup>+</sup> T cell responses for S and CD4RE pools |  |  |

**Figure 3.**
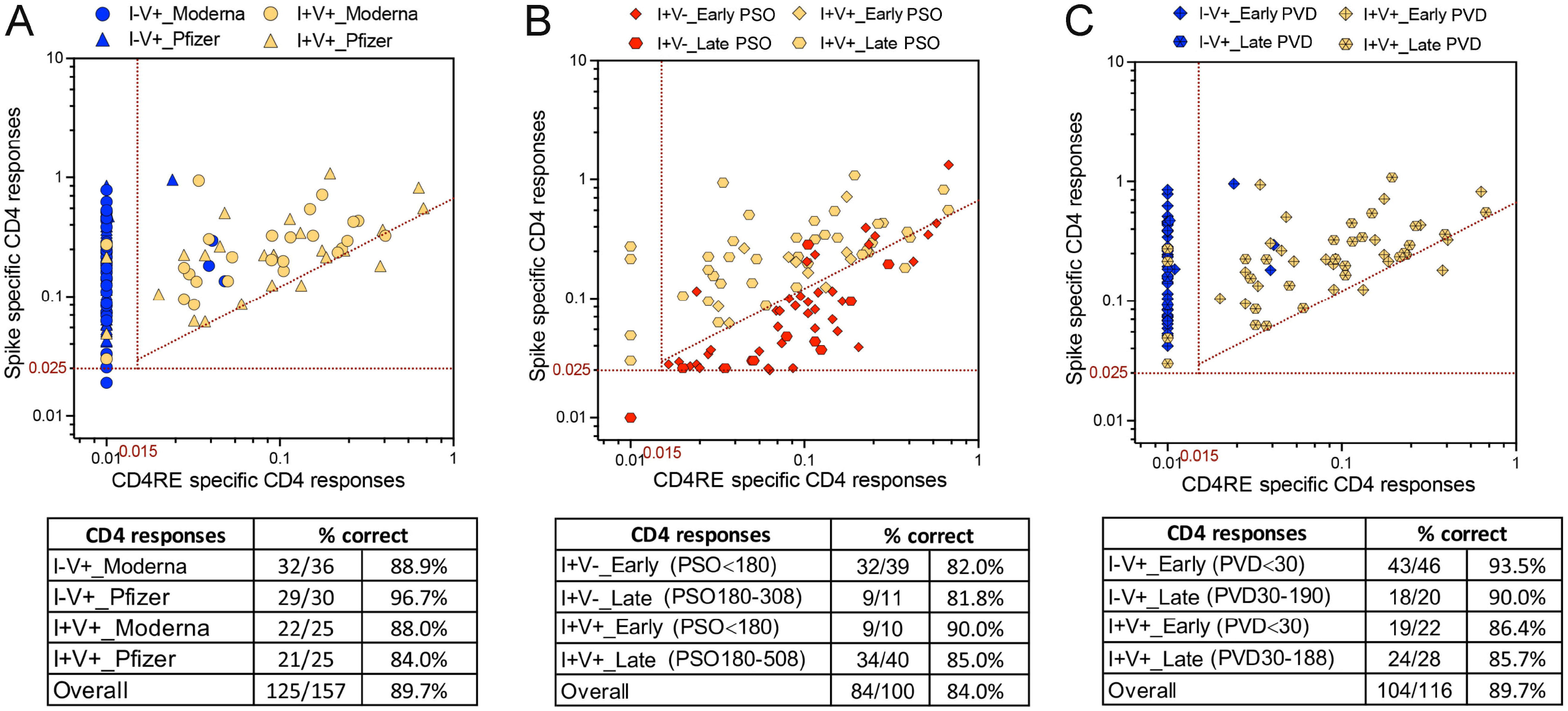
COVID-19 clinical classification scheme is applicable to different mRNA vaccines and different lengths of time post-infection/post-vaccination. CD4+ T cell responses to spike and CD4RE MPs were measured as percentage of AIM+ (OX40+CD137+) CD4+ T cells and plotted in two dimensions as absolute magnitude in order to discriminated between: (**A**) different types of mRNA vaccines (Moderna vs Pfzier) among vaccinated groups (I−V+ and I+V+); (**B**) different lengths of time post-infection among infected groups (I+V− and I+V+); (**C**) different lengths of time post-vaccination among vaccinated groups (I−V+ and I+V+). Early infection: PSO≤180; Late infection: PSO> 180; Early post-vaccination: PVD≤30; Late post-vaccination: PVD>30. I−V+, non-infected and vaccinated (n=66); I+V−, infected and non-vaccinated (n=50); I+V+, infected and then vaccinated (n=50). Red dotted lines indicate specific cutoffs. Table inserts depict the overall percentage of subjects classified correctly.

Next, we looked at the response classification as a function of the length of time PSO. The overall classification accuracy was of 84.0% (**Figure 3B**). No differences were observed in the magnitude of both total CD4+ and CD8+ T cell responses between early (≤180 days) and late (>180 days) timepoints from PSO in either the I+V− or the I+V+ groups (**Figure S5C** and **D**). The accuracy of the classification scheme when considering the different PSO timepoints was 82.0% and 81.8% in the I+V−group and 90.0% and 85.0% in the I+V+ group for the early and late timepoints, respectively (**Figure 3B**).

Lastly, we looked at the responses as a function of the length of time from the 2^nd^ dose of vaccination. The overall classification accuracy was of 89.7% (**Figure 3C**). No differences were observed in the magnitude of both total CD4+ or CD8+ T cell responses between early (≤30 days) or late (>30 days) timepoints from the last dose of vaccination in either the I−V+ or the I+V+ groups (**Figure S5E** and **F**). The accuracy of the classification scheme when considering the different vaccine timepoints was 93.5% and 90.0% in the I−V+ group and 86.4% and 85.7% in the I+V+ group for the early and late timepoints respectively. The classification scheme using CD4+ T cell AIM assay is a robust tool applicable to different types of vaccines, and can accurately classify subjects regardless of the days post-infection/post-vaccination (**Table 2**).

### CD4+ T cell reactivity of subjects associated with breakthrough infections

Breakthrough infections are defined as cases of previously COVID-19 vaccinated individuals associated with positive SARS-CoV−2 PCR tests (Bergwerk et al., 2021; Kustin et al., 2021; Mizrahi et al., 2021). Studies of antibody or T cell responses associated with breakthrough infection are scarce (Collier et al., 2021; Rovida et al., 2021). Breakthrough infection might be associated with increased immune responses as a result of the re-exposure (hybrid immunity) (Collier et al., 2021). In other cases, subjects experiencing breakthrough infections might be associated with general weaker immune responsiveness or decrease of vaccine effectiveness (Klompas, 2021; Mizrahi et al., 2021).

Here, we assessed spike and CD4RE T cell responses in a group (n=23) of breakthrough infected individuals (V+I+). Responses were compared to the vaccinated (I− V+), infected (I+V−) or infected and then vaccinated (I+V+) groups matching the V+I+ intervals of vaccination and infection (55-271 and 18-93 days, respectively). As shown in **Figure 4A**, CD4+ T cell responses from V+I+ subjects were associated with significant higher levels compared to I+V− (p=0.04) and I−V+ (p=2.3e-3) subjects and similar magnitude as the I+V+ subjects. CD8+ T cell responses had comparable levels across all the groups (**Figure 4B**). Similar to CD4+ T cell responses, spike RBD IgG titers from V+I+ subjects were equivalent to I+V+ subjects and significantly higher than I+V− (p=4.2e-7) and I−V+ (p=4.0e-15) subjects (**Figure 4C)**. Thus, at the population level breakthrough infections are associated with CD4+ T cell and spike IgG responses that resemble hybrid immunity.

**Figure 4.**
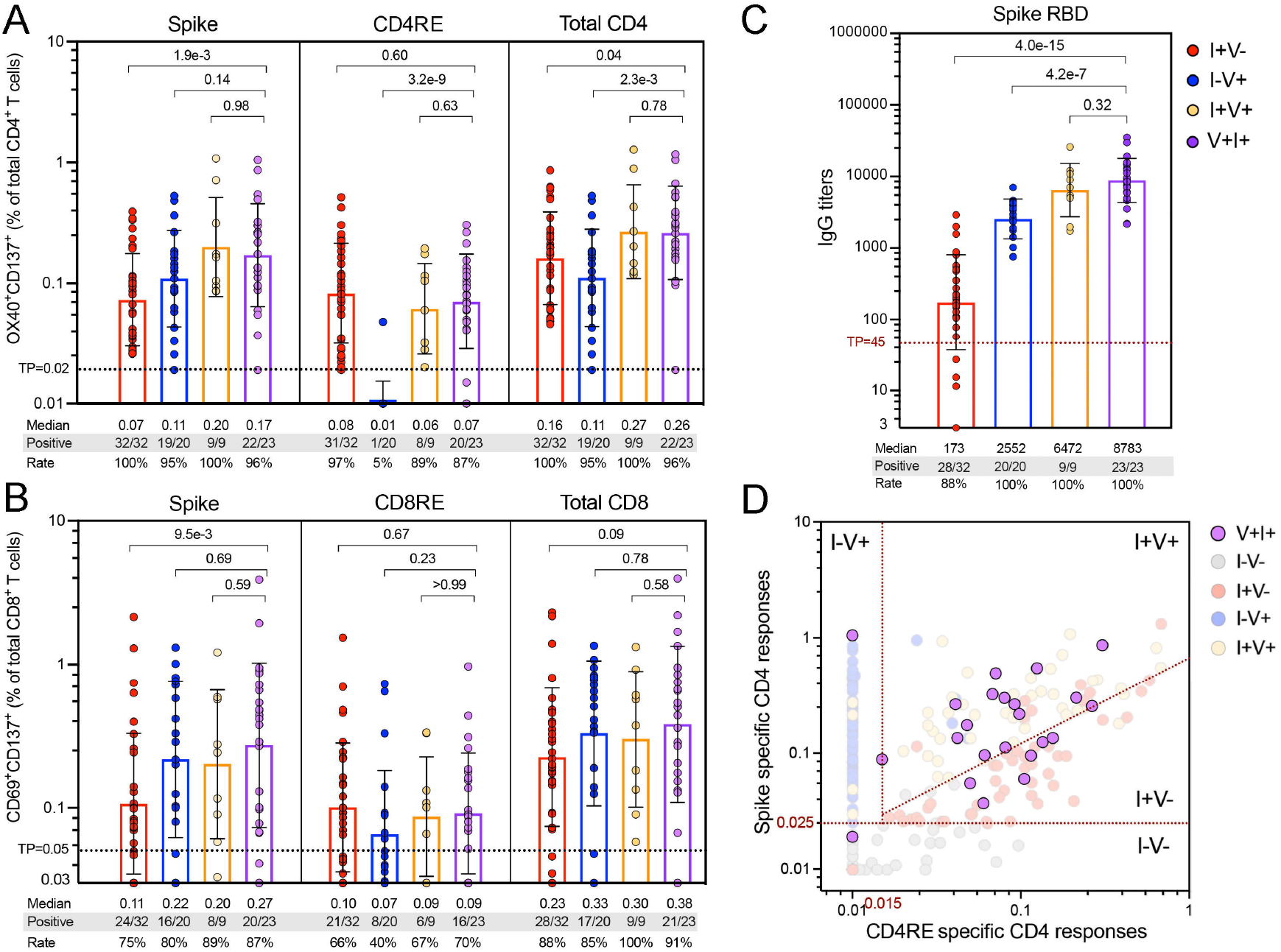
SARS-CoV−2 T cell and antibody response in breakthrough infection cases. Comparison to other study groups. SARS-CoV−2-specific T cell responses were measured as percentage of (**A**) AIM+ (OX40+CD137+) CD4+ T cells or (**B**) AIM+ (CD69+CD137+) CD8+ T cells after stimulation of PBMCs with Spike and CD4RE or CD8RE peptide pools. (**C**) Comparison of antI−spike RBD IgG titers in the plasma of the different study groups. For both T cell and antibody determinations only donors matching the V+I+ intervals of vaccination and infection (55-271 and 18-93 days, respectively) were plotted. Graph bars show geometric mean. Threshold of positivity (TP), median response, and the number or percentage of positive responding donors for each group is indicated. Kruskal-Wallis test adjusted with Dunn’s test for multiple comparisons was performed and *p* values < 0.05 considered statistically significant. (**D**) V+I+ CD4+ T cell responses plotted using the two-dimensional classification scheme with the specific cutoffs attributed to the different study groups (red dotted lines). Unexposed and unvaccinated (n=50); I+V−, infected and non-vaccinated (n=50); I+V+, infected and then vaccinated (n=50); I−V+, non-infected and vaccinated (n=66); V+I+, vaccinated and then infected (n=23).

**Figure 5.**
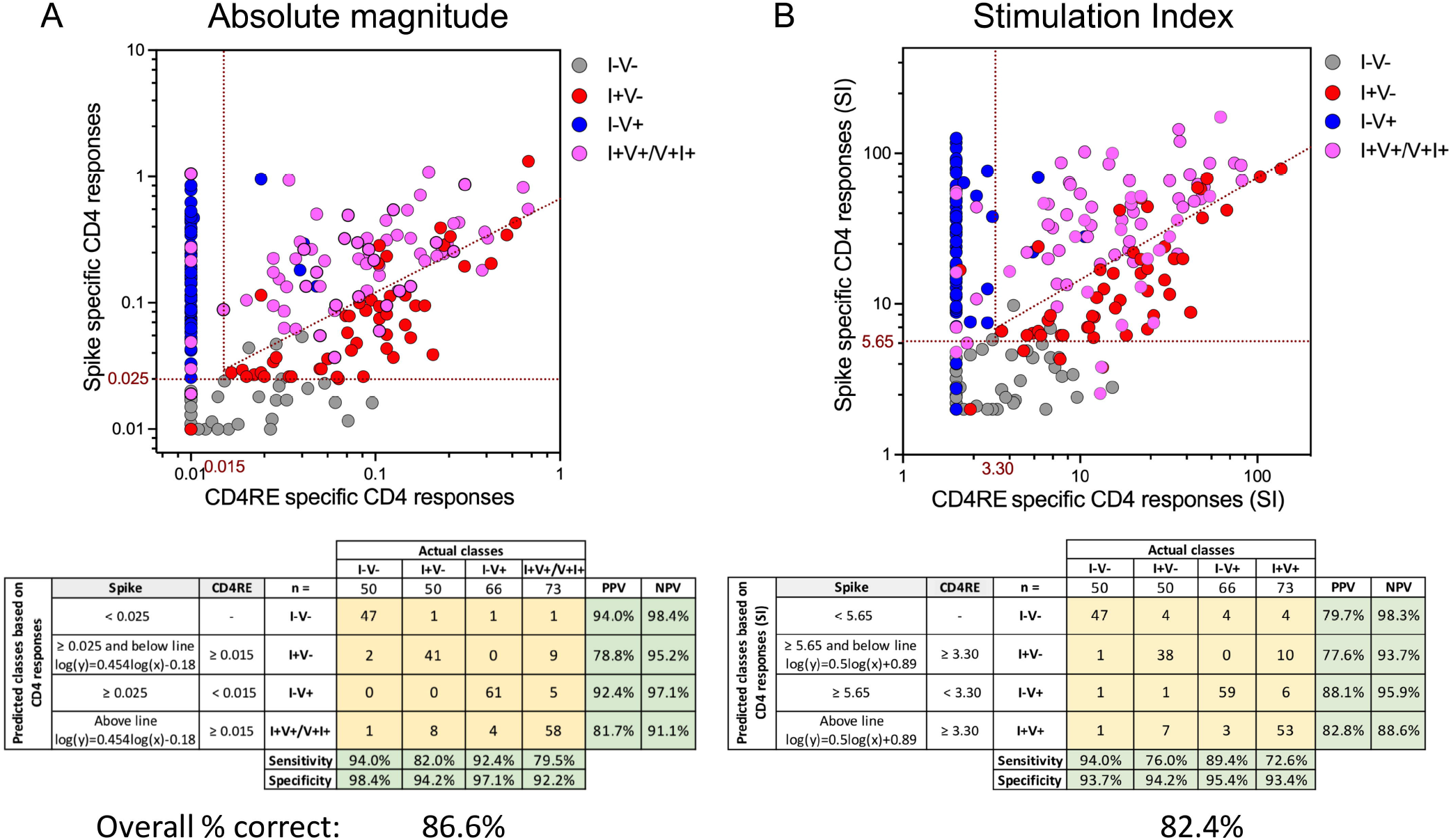
Overall COVID-19 clinical classification scheme. CD4+ T cell responses to spike and CD4RE MPs were measured as percentage of AIM+ (OX40+CD137+) CD4+ T cells and plotted in two dimensions as (**A**) SFCs per million PBMCs or (**B**) stimulation index (SI), in order to discriminate the 5 study groups with known COVID-19 status of infection, and/or vaccination. I−V−, unexposed and unvaccinated (n=50); I+V−, infected and non-vaccinated (n=50); I−V+, non-infected and vaccinated (n=66); I+V+/V+I+, infected and then vaccinated (I+V+, n=50) merged with vaccinated and then infected (V+I+, n=23). Red dotted lines indicate specific cutoffs. Table inserts depict the diagnostic exam results in 4×4 matrix. Sensitivity, specificity, PPV, NPV of all the subjects that participated in this study (n=239) and overall percentage classified correctly is shown.

### The classification scheme captures heterogeneity in breakthrough infections

At the level of the classification scheme, infections were effectively segregated from no-infected groups (unexposed and vaccinated). (**Figure 4D**). We further expected that the V+I+ breakthrough infections would be classified in the same manner of I+V+ hybrid immunity samples. Approximately two thirds (15/23 subjects) were identified by the same thresholds associated with responses from the I+V+ group (“High responders”), while the remaining third were classified similarly to I+V−subjects (“Low responders”). No obvious difference in terms of age, PSO, PVD, disease severity or length of infection from vaccination was detected between these donors and the high responders sub-group of 15 donors (**Figure S6** and **Table 1**).

In summary, while T cell responses following breakthrough infections (V+I+) are effectively segregated from the responses of uninfected donors (vaccinated or not) and follow the same pattern of responses of individuals vaccinated following natural infection (I+V+) in the majority of the cases, the classification scheme revealed heterogeneity in the CD4+ T cell responses of breakthrough donors.

### Validation of the classification scheme with whole study cohort

Finally, we summarized the overall accuracy of the classification scheme in the five cohorts used in this study including breakthrough infections. For this purpose, we clustered individuals that have been infected and vaccinated, irrespectively of the event that occurred first, into a single group, i.e. I+V/V+I+. When the 239 subjects with distinct COVID-19 status of infection and/or vaccination were combined, the classification scheme achieved a high overall accuracy, either as function of absolute magnitude (86.6%) or SI (82.4%). Also, high specificity and NPV were retained for each individual group with a range of 92.2-98.4% and 88.6-98.4% respectively. These results illustrate the highly predictive power of this classification scheme and its broad clinical applicability.

## DISCUSSION

There is a need to understand roles of SARS-CoV−2 T cell responses as potential correlates of disease outcome, and/or correlates of vaccine protection from infection or severe disease. Herein, we present the results of T cell quantitation based on the determination of relative activity directed against spike and the rest of the genome, by the use of optimized pools of experimentally defined epitopes (CD4RE and CD8RE). We report successful classification of subjects with different COVID-19 vaccination or natural infection history in the 85-90% range of accuracy. We further show that the strategy is applicable to characterizing immune responses in a group of infected vaccinees (i.e. breakthrough infections).

Although previous reports studied responses to SARS-CoV−2 in either unexposed, COVID-19 infected or vaccinated individuals (da Silva Antunes et al., 2021; Dan et al., 2021; Goel et al., 2021; Grifoni et al., 2020b; Le Bert et al., 2020; Mateus et al., 2021), this is the first demonstration, to the best of our knowledge, that a simple assay strategy can classify T-cell responses measured simultaneously in five different groups of known COVID-19 status of infection, and/or vaccination. The improved sensitivity and specificity resulted from the concept of considering the relative magnitude of responses against the spike and “rest of the genome” components, which overcomes issues related to the fact that magnitude of responses may wane over time, and also by the inclusion of experimentally defined epitopes, which we show are associated with improved signal and selectivity as compared to previously utilized predicted epitopes.

We demonstrate that the combined use of overlapping spike and CD4RE pools can be used to classify individuals with known clinical status of COVID-19, with high accuracy and sensitivity. This is of importance, as current COVID-19 diagnostic practices rely heavily on subjectively reported history, clinical records and lab modalities with imperfect performance, leading to limited reliability. For example, in longitudinal vaccination studies it will be important to monitor whether subjects enrolled in the studies might have been associated with asymptomatic infection (Kustin et al., 2021; Mizrahi et al., 2021; Pouwels et al., 2021), or even associated with abortive seronegative infections (Swadling et al., 2021). Our study supports the notion that discrimination of prior or current infections in vaccinated subjects should not rely exclusively on the analysis of spike-specific responses (Lind Enoksson et al., 2021; Martínez-Gallo, 2021; Murugesan, Preprint; Murugesan et al., 2020; Tan et al., 2021; Tormo, Pre-proof). Indeed, compared to these studies and other studies performing T cell assays using additional SARS-CoV−2 antigens (Krishna, Preprint; Kruse et al., 2021; Zelba et al., 2021), the use of our new developed pools allowed for detection of SARS-CoV−2 responses with increased sensitivity and specificity.

We also show that similar results were observed when relative versus absolute determinations were employed to measure T cell responses (i.e. using stimulation index or absolute magnitude), which allows for a more generalized use of the classification tool in different flow-cytometer platforms. The robustness of the T cell-based classification scheme was further demonstrated in an independent cohort exhibiting identical performances and was applicable to different types of mRNA vaccines, even when considering extended periods of time elapsed from infection and/or vaccination. The strength of the approach is further demonstrated by the fact that is also applicable to data generated by FluoroSpot cytokine assays despite the lower intrinsic sensitivity of this assay. We anticipate that this assay strategy will be broadly applicable to other readouts, such as ICS (Cohen et al., 2021; Mateus et al., 2021), and whole blood in an interferon-gamma release assay (IGRA) (Murugesan et al., 2020; Petrone et al., 2021; Tan et al., 2021). Although these findings were validated in several different cohorts, external validation in even larger and ethnically diverse populations and additional readouts are of interest. Potential future developments might also include epitope pools encompassing mutated epitopes from commonly circulating variants and pools with improved resolution to measure CD8+ T cell responses.

T cell responses from breakthrough infections were also evaluated. High levels of CD4+ and CD8+ T cell reactivity was observed. The elevated T cell responsiveness was paralleled by high levels of spike RBD IgG. Interestingly, these responses were of similar magnitude as responses from a group of individuals infected and then vaccinated (I+V+ in our study), whose features are commonly associated with hybrid immunity (Crotty, 2021). Notably, breakthrough infections were also associated with higher CD4+ T cell and spike RBD IgG responses compared to infected only or vaccinated only subjects. These results suggest that T and B cell reactivity associated with breakthrough infections is increased as a result of re-exposure. However, the classification tool system, also revealed significant heterogeneity in responses in some subjects, possibly linking breakthrough infections to lower adaptive responses.

## ACKNOWLEDGEMENTS

We wish to acknowledge all subjects for their participation and for donating their blood and time for this study. We are grateful to the La Jolla Institute for Immunology clinical core relentless efforts and for the support of Sanguine, BioIVT, and iSpecimen in obtaining blood samples. Research reported in this publication was supported by the National Institute of Allergy and Infectious Diseases (NIAID) of the National Institutes of Health (NIH) under Award number U19 AI142742, and contract number 75N93019C00065 and 5N9301900066. The content is solely the responsibility of the authors and does not necessarily represent the official views of the National Institutes of Health.

## AUTHOR CONTRIBUTIONS

Conceptualization: E.D.Y., R.d.S.A., and A.Se. Methodology: E.D.Y., J.M.D., E.W., E.G., A.Su., B.G., J.C., R.I.G., J.M., and A.G. Formal analysis: E.D.Y., J.M.D., R.d.S.A, and A.Se. Investigation: E.D.Y., S.C., R.d.S.A. and A.Se. Funding acquisition: D.W., S.C., and A.Se. Writing: E.D.Y., S.C., R.d.S.A., and A.Se. Supervision: E.D.Y., R.d.S.A., S.C., and A.Se.

## DECLARATION OF INTERESTS

A.Se. is a consultant for Gritstone Bio, Flow Pharma, Arcturus Therapeutics, ImmunoScape, CellCarta, Avalia, Moderna, Fortress and Repertoire. S.C. is a consultant for Avalia. LJI has filed for patent protection for various aspects of SARS-CoV−2 epitope pools design. All other authors declare no conflict of interest.

## MATERIAL AND METHODS

### LEAD CONTACT AND MATERIALS AVAILABILITY

Please contact A.S. and R.d.S.A for aliquots of synthesized sets of peptides identified in this study. There are restrictions to the availability of the peptide reagents due to cost and limited quantity.

### Human Subjects and PBMC isolation

The Institutional Review Boards of the University of California, San Diego (UCSD; 200236X) and the La Jolla Institute for Immunology (LJI; VD-214) approved the protocols used for blood collection for all the subjects who donated at all sites. The vast majority of the blood donations were collected through the UC San Diego Health Clinic and at the La Jolla Institute for Immunology (LJI). Additional samples were obtained from contract research organizations (CRO) under the same LJI IRB approval. All samples with the exception of the I−V−study group were collected during COVID-19 pandemic from 2020-2021. Pre-pandemic blood donations of the I−V−group were performed from 2013-2019. Each participant provided informed consent and was assigned a study identification number with clinical information recorded. Subjects who had a medical history and/or symptoms consistent with COVID-19, but lacked positive PCR-based testing for SARS-CoV−2 and subsequently had negative laboratory-based serologic testing for SARS-CoV−2, were then excluded; i.e., all COVID-19 cases in this study were confirmed cases by SARS-CoV−2 PCR or SARS-CoV−2 serodiagnostics, or both. Adults of all races, ethnicities, ages, and genders were eligible to participate. Study exclusion criteria included lack of willingness to participate, lack of ability to provide informed consent, or a medical contraindication to blood donation (e.g., severe anemia). In all cases, PBMCs were isolated from whole blood by density gradient centrifugation according to manufacturer instructions (Ficoll-Hypaque, Amersham Biosciences, Uppsala, Sweden). Cells were cryopreserved in liquid nitrogen suspended in FBS containing 10% (vol/vol) DMSO (Sigma-Aldrich). Plasma was obtained by centrifugation (400g for 15 minutes at 4°C) of whole blood and collection of the upper layer, prior to PBMC isolation and cryopreserved at −80°C.

### Design and production of new SARS-CoV−2 epitope pools

To study T cell responses against SARS-CoV−2, we used a megapool (MP) of 15-mer peptides overlapping by 10 spanning the entire spike protein sequence (253 peptides; **Table S3**) as previously described (Grifoni et al., 2020b). For the rest of the SARS-CoV−2 proteome, and in order to design epitope pools with increased HLA coverage and broadly recognized by demographically and geographically diverse populations, experimental defined epitopes from non-spike (R) region of SARS-CoV−2 were selected based on our recent meta-analysis (Grifoni et al., 2021). Briefly, peptides were synthetized and pooled to include both dominant (recognized in 3 or more donors/studies) and subdominant epitopes. To improve specificity, overly short or long ligands which could cause “false positive” signals (Paul et al., 2018), were excluded and only peptides of sizes ranging 15-20 and 9-10 amino acids, respectively in CD4RE and CD8RE pools were included, resulting in the generation of CD4RE and CD8RE MPs with 284 and 621 peptides, respectively (**Table S3**). Detailed information of the MPs composition with peptide sequences, length, ORFs of origin, and HLA coverages are specified in **Table S4**. Alternatively, a MP for the remainder genome consisting of dominant HLA class II predicted CD4+ T-cell epitopes (221 peptides), as previously described (Grifoni et al., 2020b) was also used as control (**Table S3)**. In addition, an EBV pool of previously reported experimental class I and class II epitopes (Carrasco Pro et al., 2015) with 301 peptides was used as positive control. All peptides were synthesized by TC peptide lab (San Diego, CA), pooled and resuspended at a final concentration of 1□mg/mL in DMSO.

### SARS-CoV−2 RBD Spike and Nucleocapsid ELISAs

The SARS-CoV−2 ELISAs have been described in detail previously (Dan et al., 2021). Briefly, 96-well half-area plates (ThermoFisher 3690) were coated with 1 ug/mL of antigen and incubated at 4°C overnight. Antigens included recombinant SARS-CoV−2 RBD protein obtained from the Saphire laboratory at LJI or recombinant nucleocapsid protein (GenScript Z03488). The next day, plates were blocked with 3% milk in phosphate-buffered saline (PBS) containing 0.05% Tween-20 for 1.5 hours at room temperature. Plasma was heat inactivated at 56°C for 30 to 60 min. Plasma was diluted in 1% milk containing 0.05% Tween-20 in PBS starting at a 1:3 dilution followed by serial dilutions by three and incubated for 1.5 hours at room temperature. Plates were washed five times with 0.05% PBS-Tween-20. Secondary antibodies were diluted in 1% milk containing 0.05% Tween-20 in PBS. AntI−human IgG peroxidase antibody produced in goat (Sigma A6029) was used at a 1:5,000 dilution. Subsequently, plates were read on Spectramax Plate Reader at 450 nm, and data analysis was performed using SoftMax Pro. End-point titers were plotted for each sample, using background-subtracted data. Negative and positive controls were used to standardize each assay and normalize across experiments. Limit of detection (LOD) was defined as 1:3 of IgG. Spike RBD IgG or nucleocapsid IgG thresholds of positivity (TP) for SARS-CoV−2 infected or COVID-19 vaccinated individuals were established based on uninfected and unvaccinated subjects (I−V−).

### Activation induced cell marker (AIM) assay

The AIM assay was performed as previously described (Mateus et al., 2020). Cryopreserved PBMCs were thawed by diluting the cells in 10 mL complete RPMI 1640 with 5% human AB serum (Gemini Bioproducts) in the presence of benzonase [20 ml/10ml]. Cells were cultured for 20 to 24 hours in the presence of SARS-CoV−2 specific and EBV pools (1ug/ml) in 96-wells U bottom plates with 1×10^6^ PBMC per well. An equimolar amount of DMSO was added as a negative control and phytohemagglutinin (PHA, Roche (San Diego, CA) 1 mg/ml) was used as the positive control. The cells were stained with CD3 AF532, CD4 BV605, CD8 BUV496, and Live/Dead Aqua. Activation was measured by the following markers: CD137 APC, OX40 PE-Cy7, and CD69 PE. The detailed information of the antibodies used are summarized in **Table S5**. All samples were acquired on a ZE5 cell analyzer (Biorad laboratories, Hercules, CA) and analyzed with FlowJo software (Tree Star, Ashland, OR).

CD4+ and CD8+ T cells responses were calculated as percent of total CD4+ (OX40^+^CD137^+^) or CD8+ (CD69^+^CD137^+^) T cells. The background was removed from the data by subtracting the wells stimulated with DMSO. The Stimulation Index (SI) was calculated by dividing the counts of AIM+ cells after SARS-CoV−2 pools stimulation with the ones in the negative control. A positive response was defined as SI≥2 and AIM^+^ response above the threshold of positivity after background subtraction. The limit of detection (0.01% and 0.03 for CD4+ and CD8+ T cells, respectively) was calculated based on 2 times 95% CI of geomean of negative control (DMSO), and the threshold of positivity (0.02% for CD4+ and 0.05% for CD8+ T cells) was calculated based on 2 times standard deviation of background signals according to previous published studies (Dan et al., 2021; Mateus et al., 2020). The gating strategy utilized is shown in **Figure S7**, as well as reactive CD4+ and CD8+ T cell responses to SARS-CoV−2, EBV and PHA positive control from a representative donor.

### IFNγ FluoroSpot assay

The FluoroSpot assay was performed as previously described (Tarke et al., 2021a). PBMCs derived from 80 subjects from 4 clinical cohorts (20 each for I−V−, I+V−, I−V+, and I+V+ cohorts) were stimulated in triplicate at a single density of 2×10^5^ cells/well. The cells were stimulated with the different MPs analyzed (1ug/mL), PHA (10mg/mL), and DMSO (0.1%) in 96-well plates previously coated with antI−cytokine antibodies for IFNγ, (mAbs 1-D1K; Mabtech, Stockholm, Sweden) at a concentration of 10ug/mL. After 20-24 hours of incubation at 37°C, 5% CO2, cells were discarded and FluoroSpot plates were washed and further incubated for 2 hours with cytokine antibodies (mAbs 7-B6-1-BAM; Mabtech, Stockholm, Sweden). Subsequently, plates were washed again with PBS/0.05% Tween20 and incubated for 1 hour with fluorophore-conjugated antibodies (AntI−BAM-490). Computer-assisted image analysis was performed by counting fluorescent spots using an AID iSPOT FluoroSpot reader (AIS-diagnostika, Germany). Each megapool was considered positive compared to the background based on the following criteria: 20 or more spot forming cells (SFC) per 10^6^ PBMC after background subtraction for each cytokine analyzed, a stimulation index (SI) greater than 2, and statistically different from the background (p < 0.05) in either a Poisson or t test.

### Statistical analysis

Experimental data were analyzed by GraphPad Prism Version 9 (La Jolla, CA) and Microsoft Excel Version 16.16.27 (Microsoft, Redmond, WA). The statistical details of the experiments are provided in the respective figure legends. Data were analyzed by Wilcoxon test (two-tailed) to compare between two paired groups, and Kruskal-Wallis test adjusted with Dunn’s test for multiple comparisons to compare between multiple groups. Data were plotted as geometric mean with geometric SD. *p* values < 0.05 (after adjustment if indicated) were considered statistically significant. For the classification scheme, statistical determinations and metrics were executed as previously described (Trevethan, 2017). Briefly, for each individual group the following calculations were performed: 1) positive predictive value (PPV)= (True Positives)/(True Positives+False Positives); 2) negative predictive value (NPV)= (True Negatives)/(True Negatives+False Negatives); 3) sensitivity =(True Positives)/(True Positives+False Negatives); and 4) specificity=(True Negatives)/(True Negatives+False Positives).

## STUDY APPROVAL

This study was approved by the Human Subjects Protection Program of the UC San Diego Health under IRB approved protocols (UCSD; 200236X), or under IRB approval (LJI; VD-214) at the La Jolla Institute for Immunology. All donors were able to provide informed consent, or had a legal guardian or representative able to do so. Each participant provided informed consent and was assigned a study identification number with clinical information recorded.

## DATA AVAILABILITY

The datasets generated and analyzed in this study are available from the corresponding authors upon reasonable request. Likewise, biomaterials archived from this study may be shared for further research.

## SUPPLEMENTAL INFORMATION

Supplemental Information can be found in attached file: Supplemental figures and tables.pdf.

