## Supplemental Figures and Tables for "Distinguishing COVID-19 infection and vaccination history by T cell reactivity"

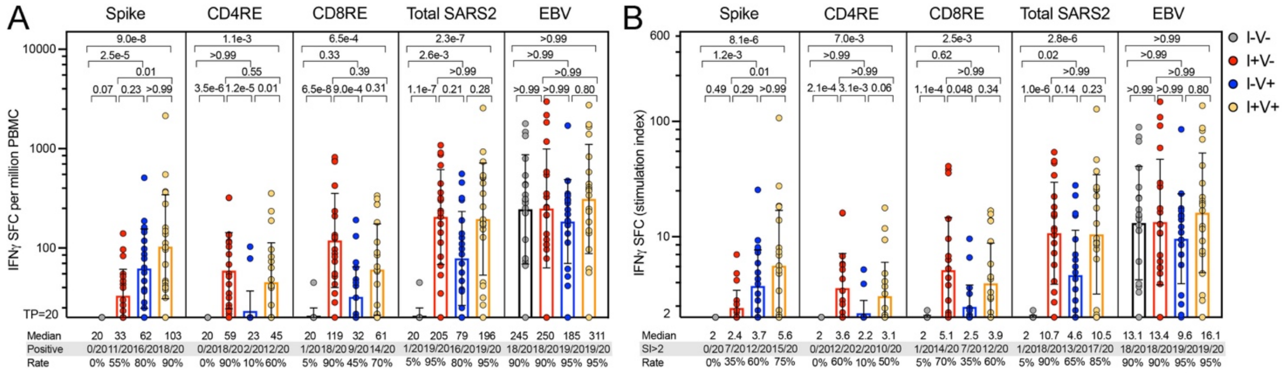


**Figure S1. SARS-CoV-2-specific CD4+ and CD8+ T cell IFNγ responses assessed by FluoroSpot**

IFNγ production was measured by FluoroSpot after stimulation of PBMCs with peptides pools encompassing spike only (Spike) MP or the experimentally defined CD4RE and CD8RE MPs representing all the proteome without spike. EVB MP was used as a control. Graphs show individual response of spike, CD4RE, CD8RE or the combination of all (Total SARS2) plotted as background subtracted (**A**) or as SI (**B**) against DMSO negative control. Geometric mean for the 4 different groups is shown. Kruskal-Wallis test adjusted with Dunn's test for multiple comparisons. Geometric mean for the 4 different groups is shown. Kruskal-Wallis test adjusted with Dunn's test for multiple comparisons was performed and *p* values < 0.05 considered statistically significant. I-V-, unexposed and unvaccinated (n=20); I+V-, infected and non-vaccinated (n=20); I+V+, infected and then vaccinated (n=20); I-V+, non-infected and vaccinated (n=20). Threshold of positivity (TP) is indicated. Median response, and the number or percentage of positive responding donors for each group is shown.

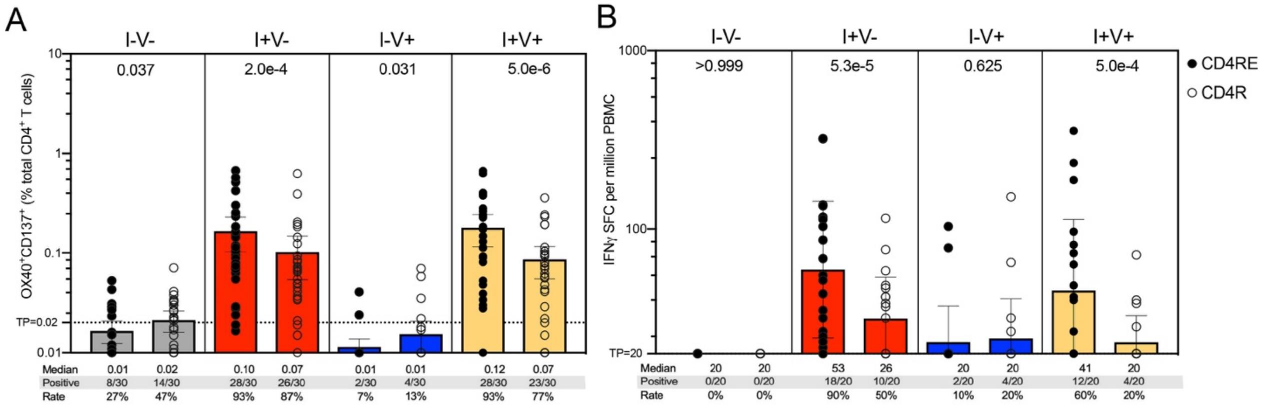


**Figure S2. Comparison between CD4RE and CD4R MP performances in the detection of SARS-CoV-2 CD4+ T cell responses measured in activation induced marker (AIM) and FluoroSpot assays**

SARS-CoV-2 experimentally defined CD4RE or predicted CD4R MPs were used to stimulate PBMCs and reactive CD4+ T cell responses measured by both AIM and FluoroSpot assays. (**A**) Percentage of AIM+ (OX40+CD137+) CD4+ T cells. (**B**) IFNγ production represented as SFCs per million PBMCs. Bars represent geometric mean ± geometric SD. *p* values were calculated by Wilcoxon test (two-tailed), and $p < 0.05$ considered statistically significant. I-V-, unexposed and unvaccinated; I+V-, infected and non-vaccinated; I+V+, infected and then vaccinated; I-V+, non-infected and vaccinated. n=30 or n=20 were included in each group for AIM or FluoroSpot assays, respectively. Threshold of positivity (TP) is indicated. Median response, and the number or percentage of positive responding donors for each group is shown.

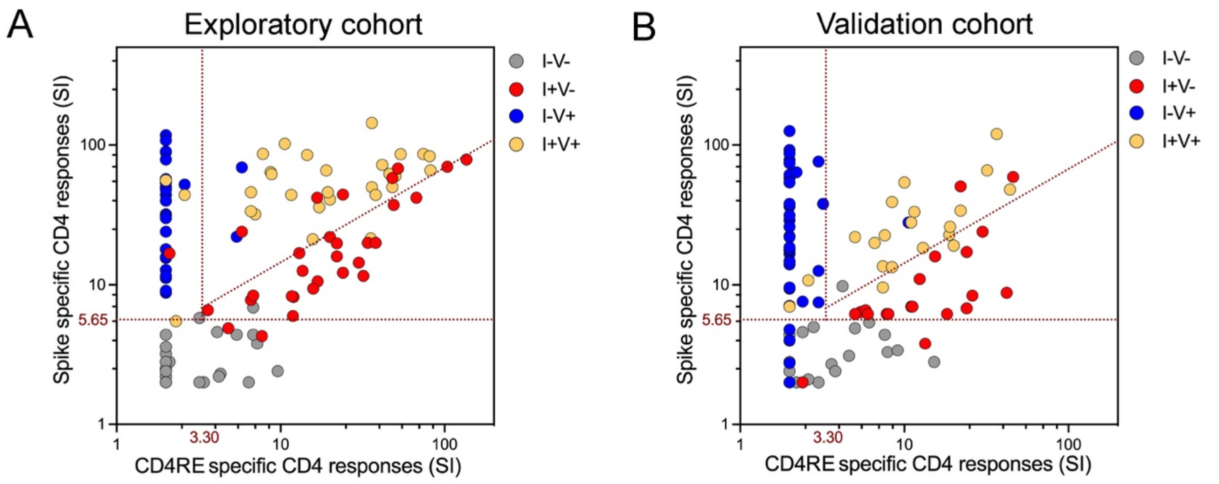


| Predicted classes based on CD4 responses (SI): Spike | CD4RE | n = | Actual classes: I-V- (30) | I+V- (30) | I-V+ (30) | I+V+ (30) | PPV | NPV |
|---|---|---|---|---|---|---|---|---|
| < 5.65 | - | I-V- | 28 | 2 | 0 | 1 | 90.3% | 97.8% |
| ≥ 5.65 and below line log(y)=0.5log(x)+0.89 | ≥ 3.30 | I+V- | 1 | 22 | 0 | 1 | 91.7% | 91.7% |
| ≥ 5.65 | < 3.30 | I-V+ | 1 | 1 | 28 | 2 | 87.5% | 97.7% |
| Above line log(y)=0.5log(x)+0.89 | ≥ 3.30 | I+V+ | 0 | 5 | 2 | 26 | 78.8% | 95.4% |
| | | Sensitivity | 93.3% | 73.3% | 93.3% | 86.7% | | |
| | | Specificity | 96.7% | 97.8% | 95.6% | 92.2% | | |

| Predicted classes based on CD4 responses (SI): Spike | CD4RE | n = | Actual classes: I-V- (20) | I+V- (20) | I-V+ (36) | I+V+ (20) | PPV | NPV |
|---|---|---|---|---|---|---|---|---|
| < 5.65 | - | I-V- | 19 | 2 | 4 | 0 | 76.0% | 98.6% |
| ≥ 5.65 and below line log(y)=0.5log(x)+0.89 | ≥ 3.30 | I+V- | 0 | 16 | 0 | 2 | 88.9% | 94.9% |
| ≥ 5.65 | < 3.30 | I-V+ | 0 | 0 | 31 | 2 | 93.9% | 92.1% |
| Above line log(y)=0.5log(x)+0.89 | ≥ 3.30 | I+V+ | 1 | 2 | 1 | 16 | 80.0% | 94.7% |
| | | Sensitivity | 95.0% | 80.0% | 86.1% | 80.0% | | |
| | | Specificity | 92.1% | 97.4% | 96.7% | 94.7% | | |

Overall % correct: 86.7% 85.4%

**Figure S3. COVID-19 clinical classification scheme can classify subjects from I-V-, I+V-, I-V+, and I+V+ cohorts using stimulation index readouts**

Related to Figure 2, CD4+ T cell responses to spike and CD4RE MPs were measured as percentage of AIM+ (OX40+CD137+) CD4+ T cells and plotted as stimulation index (SI) in two dimensions in order to discriminate the 4 study groups with known COVID-19 status of infection, and/or vaccination in 2 independent cohorts: (**A**) Exploratory cohort (n=120) and (**B**) Validation cohort (n=96). I-V-, unexposed and unvaccinated (n=30 and n=20); I+V-, infected and non-vaccinated (n=30 and n=20); I+V+, infected and then vaccinated (n=30 and n=20); I-V+, non-infected and vaccinated (n=30 and n=36). Red dotted lines indicate specific cutoffs. Table inserts depict the diagnostic exam results in 4x4 matrix. Sensitivity, specificity, PPV, NPV and overall percentage of subjects classified correctly is shown.

A

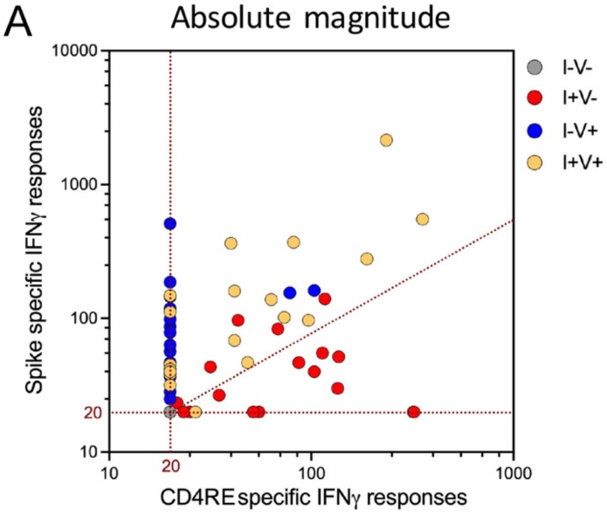


| Predicted classes based on IFNγ responses: Spike | CD4RE | n = | Actual classes: I-V- (20) | I+V- (20) | I-V+ (20) | I+V+ (20) | PPV | NPV |
|---|---|---|---|---|---|---|---|---|
| ≤ 20 | ≤ 20 | I-V- | 20 | 2 | 4 | 1 | 74.1% | 100.0% |
| ≥ 20 and below line log(y)=0.84log(x)+0.2 | > 20 | I+V- | 0 | 13 | 0 | 1 | 92.9% | 89.4% |
| > 20 | ≤ 20 | I-V+ | 0 | 0 | 14 | 7 | 66.7% | 89.8% |
| Above line log(y)=0.84log(x)+0.2 | > 20 | I+V+ | 0 | 5 | 2 | 11 | 61.1% | 85.5% |
| | | Sensitivity | 100.0% | 65.0% | 70.0% | 55.0% | | |
| | | Specificity | 88.3% | 98.3% | 88.3% | 88.3% | | |

Overall accuracy: 72.5%

B

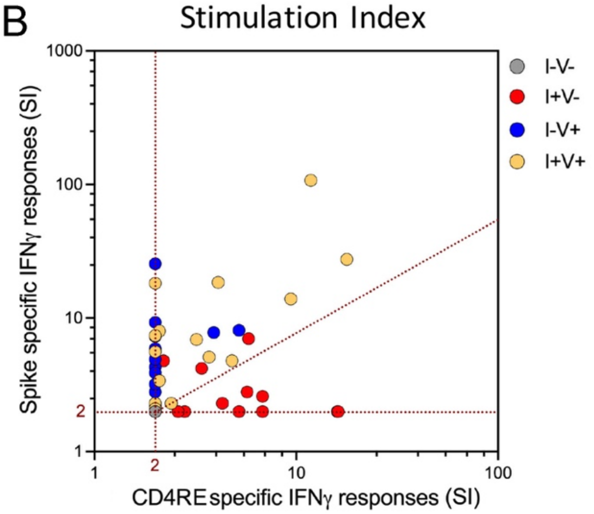


| Predicted classes based on IFNγ responses (SI): Spike | CD4RE | n = | Actual classes: I-V- (20) | I+V- (20) | I-V+ (20) | I+V+ (20) | PPV | NPV |
|---|---|---|---|---|---|---|---|---|
| ≤ 2 | ≤ 2 | I-V- | 20 | 7 | 8 | 5 | 50.0% | 100.0% |
| ≥ 2 and below line log(y)=0.84log(x)+0.2 | > 2 | I+V- | 0 | 9 | 0 | 1 | 90.0% | 84.3% |
| > 2 | ≤ 2 | I-V+ | 0 | 1 | 10 | 5 | 62.5% | 84.4% |
| Above line log(y)=0.84log(x)+0.2 | > 2 | I+V+ | 0 | 3 | 2 | 9 | 64.3% | 83.3% |
| | | Sensitivity | 100.0% | 45.0% | 50.0% | 45.0% | | |
| | | Specificity | 66.7% | 98.3% | 90.0% | 91.7% | | |

60.0%

**Figure S4. COVID-19 clinical classification scheme using IFNγ responses assessed by FluoroSpot. Related to Figure 2**

IFNγ production to spike and CD4RE MPs were measured by FluoroSpot and responses plotted in two dimensions as (**A**) SFCs per million PBMCs or (**B**) stimulation index (SI) in order to discriminate the 4 study groups with known COVID-19 status of infection, and/or vaccination. I-V-, unexposed and unvaccinated (n=20); I+V-, infected and non-vaccinated (n=20); I+V+, infected and then vaccinated (n=20); I-V+, non-infected and vaccinated (n=20). Red dotted lines indicate specific cutoffs. Table inserts depict the diagnostic exam results in 4x4 matrix. Sensitivity, specificity, PPV, NPV and overall percentage of subjects classified correctly is shown.

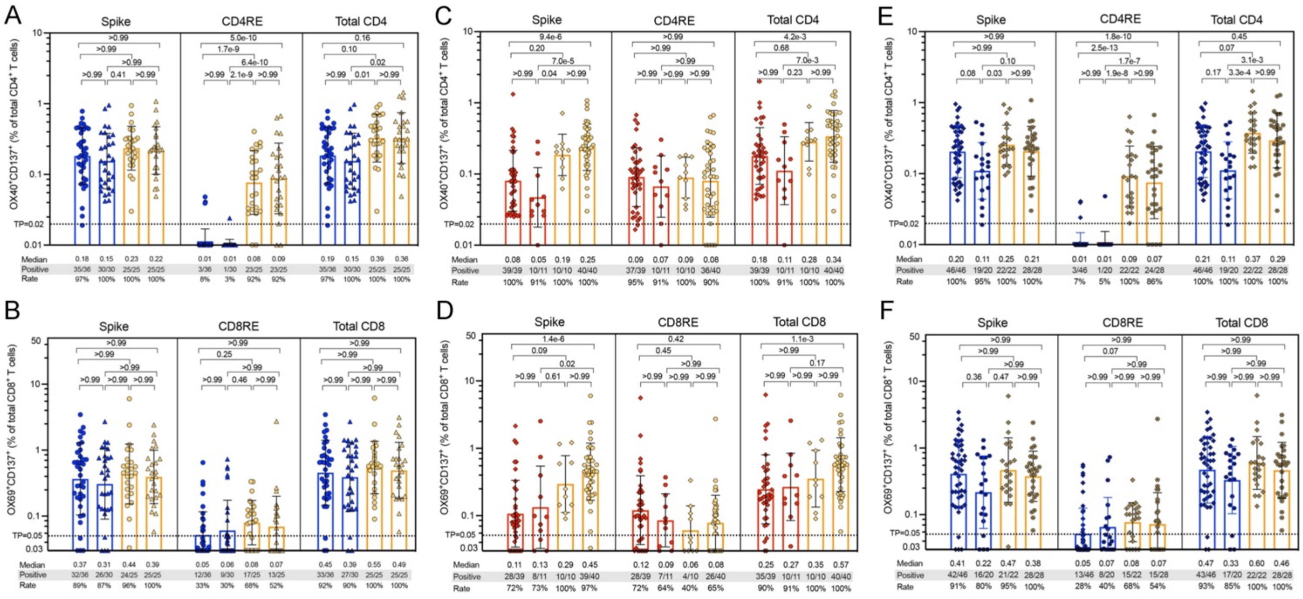


**Figure S5. SARS-CoV-2-specific CD4+ and CD8+ T cell responses in different mRNA vaccines and different lengths of time post-infection/post-vaccination**

SARS-CoV-2-specific T cell responses were measured as percentage of AIM+ (OX40+CD137+) CD4+ T cells or AIM+ (CD69+CD137+) CD8+ T cells after stimulation of PBMCs with peptides pools encompassing spike only (Spike) MP or the experimentally defined CD4RE and CD8RE MPs representing all the proteome without spike. Graphs show individual response of spike, CD4RE or CD8RE and the combination of both (Total CD4+ or Total CD8+) plotted as background subtracted for (**A, B**) different types of mRNA vaccines (Moderna vs Pfzier) among vaccinated groups (I-V+ and I+V+); (**C, D**) different lengths of time post-infection among infected groups (I+V- and I+V+); (**E, F**) different lengths of time post-vaccination among vaccinated groups (I-V+ and I+V+). Early infection: PSO≤180; Late infection: PSO> 180; Early post-vaccination: PVD≤30; Late post-vaccination: PVD>30. I-V+, non-infected and vaccinated (n=66); I+V-, infected and non-vaccinated (n=50); I+V+, infected and then vaccinated (n=50). Geometric mean for the 4 different groups is shown. Kruskal-Wallis test adjusted with Dunn's test for multiple comparisons was performed and *p* values < 0.05 considered statistically significant. Threshold of positivity (TP) is indicated. Median response, and the number or percentage of positive responding donors for each group is shown.

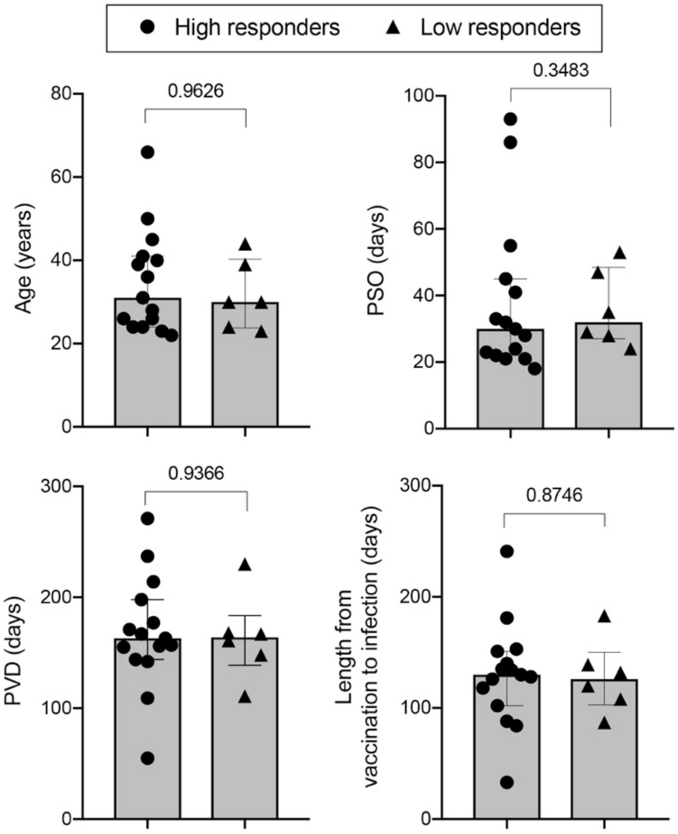


**Figure S6. Individuals with high breakthrough infection responses have similar clinical characteristics to individuals with low responses**

Breakthrough infected subjects were either grouped as “high responders” (n=15; identified by the same thresholds associated with responses from the I+V+ group), or as “low responders” (n=6; identified by the same thresholds associated with responses from the I+V- group) and plotted for age, PSO, PVD or length of infection from vaccination. High and Low responders are represented by circles and triangles, respectively. Median with interquartile range is shown and *p* values are indicated.

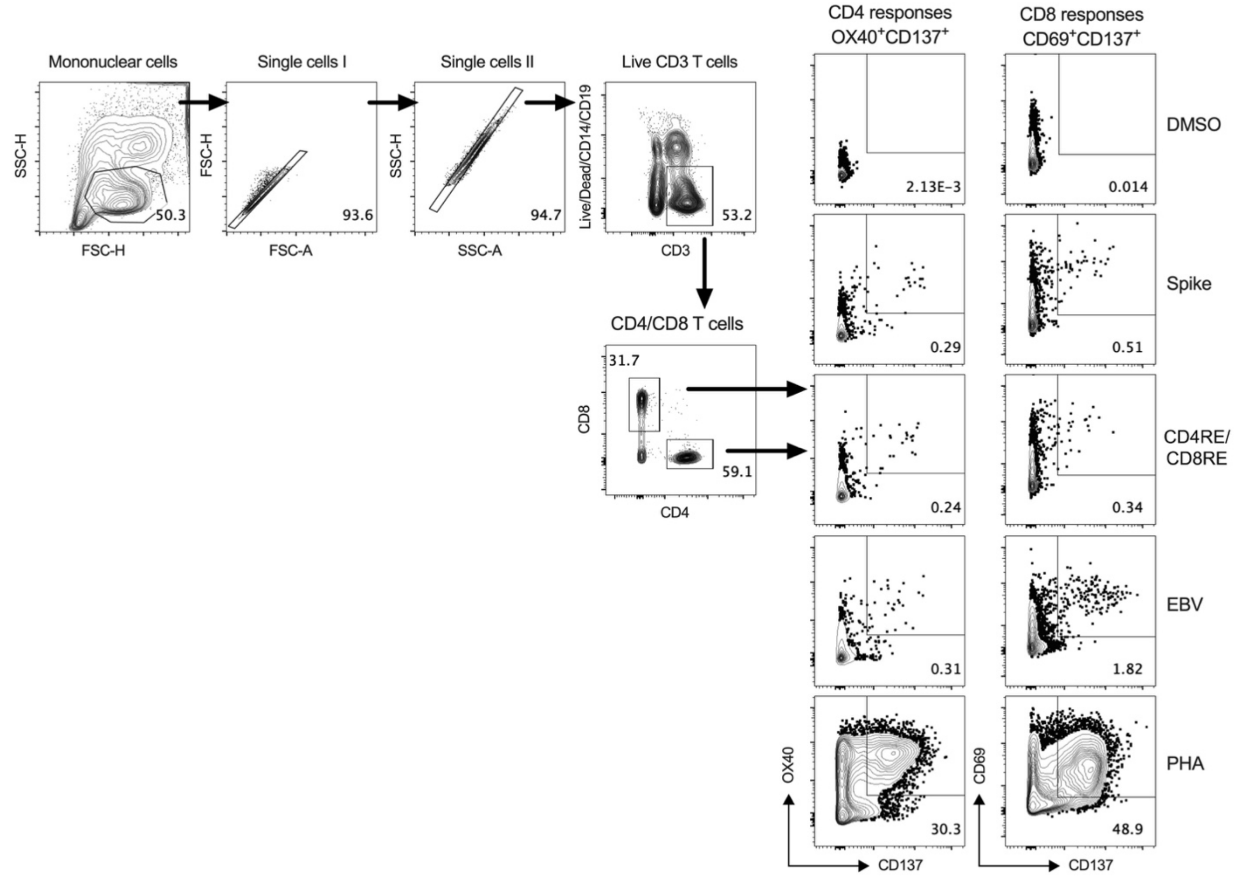


**Figure S7. Gating strategy and representative CD4 and CD8 responses plots of the activation induced marker (AIM) assay**

Representative gating of live CD3+ T cells, CD4+ T cells, CD8+ T cells and reactive OX40+CD137+ CD4 or CD69+CD137+ CD8 T cells from donor PBMCs is shown. Briefly, mononuclear cells were gated out of all events followed by subsequent singlet gating. Live CD3+ cells were gated as Live/Dead-CD14-CD19-CD3+. Cells were then gated as CD4+CD8- or CD4-CD8+ T cells, and reactive OX40+CD137+ CD4 or CD69+CD137+ CD8 T cells were gated and calculated as percent of total CD4 or CD8 T cells. Representative CD4 and CD8 responses plots after stimulating with positive (PHA and EBV) or negative (DMSO) controls and SARS-CoV-2 specific megapools were presented on the right side.

**Table S1. Description of donor cohort characteristics and demographics for the experimental cohort**

| Study cohort | I-V- group | I+V- group | I-V+ group | I+V+ group |
|---|---|---|---|---|
| Number of subjects | 30 | 30 | 30 | 30 |
| Age | 24 (17-64) | 42 (20-67) | 41 (24-69) | 43 (21-73) |
| Gender (female%) | 53% | 70% | 67% | 43% |
| Ethnicity | | | | |
| Caucasian | 19 | 23 | 16 | 25 |
| Hispanic/Latino | 5 | 4 | 6 | 2 |
| Asian | 5 | 2 | 7 | 2 |
| African American | 1 | 1 | 1 | 1 |
| Infection confirmed by PCR test | - | Yes | - | Yes |
| SARS-CoV-2 S protein RBD IgG titers | 3.0 (3.0-3.0) | 182.3 (3.0-7326.5) | 3932.6 (410.6-13677.0) | 3609.6 (159.2-12060.6) |
| Days from symptom onset | - | 100 (20-290) | - | 380 (57-460) |
| Vaccine type | | | | |
| BNT162b2 mRNA (Pfizer/BioNTech) | - | - | 15 (50%) | 15 (50%) |
| mRNA-1273 (Moderna) | - | - | 15 (50%) | 15 (50%) |
| Days from 2$^{nd}$ dose of vaccination | - | - | 14 (13-77) | 30 (7-85) |

**Table S2. Description of donor cohort characteristics and demographics for the validation cohort**

| Validation cohort | I-V- group | I+V- group | I-V+ group | I+V+ group |
|---|---|---|---|---|
| Number of subjects | 20 | 20 | 36 | 20 |
| Age | 26 (18-37) | 40 (19-67) | 40 (21-74) | 33 (21-65) |
| Gender (female%) | 40% | 40% | 50% | 70% |
| Ethnicity | | | | |
| Caucasian | 13 | 14 | 22 | 14 |
| Hispanic/Latino | 3 | 3 | 5 | 4 |
| Asian | 3 | 2 | 9 | 1 |
| African American | 1 | 1 | 1 | 1 |
| Infection confirmed by PCR test | - | Yes | - | Yes |
| SARS-CoV-2 S protein RBD IgG titers | 4.5 (3.0-23.9) | 207.3 (5.2-2904.8) | 4372.5 (755.0-32033.0) | 7343.0 (564.6-25876.0) |
| Days from symptom onset | - | 162 (39-308) | - | 265 (120-508) |
| Vaccine type | | | | |
| BNT162b2 mRNA (Pfizer/BioNTech) | - | - | 15 (42%) | 10 (50%) |
| mRNA-1273 (Moderna) | - | - | 21 (58%) | 10 (50%) |
| Days from 2$^{nd}$ dose of vaccination | - | - | 45 (13-190) | 72 (13-188) |

### Table S3. Megapools used to detect CD4+ and CD8+ T cell responses

| Name | Source | Peptide/Epitope type | Number of peptides | Reference |
|---|---|---|---|---|
| Spike | Spike | Overlapping | 253 | Grifoni et al., Cell (2020) |
| CD4R | Non-spike proteome | Predicted | 221 | |
| CD4RE | Non-spike proteome | Exp defined dominant and subdominant | 284 | Grifoni et al., Cell Host Microbe (2021) |
| CD8RE | Non-spike proteome | Exp defined dominant and subdominant | 621 | |
| EBV | Whole proteome | Exp defined | 301 | IEDB |

### Table S4. Detailed peptide sequences information of CD4RE and CD8RE MP

Please see attached excel file: Supplementary Table 4.xlsx

### Table S5. Detailed information of the antibodies used in this study

| Antibody | Fluorochrome | Clone | Vendor | Catalog number |
|---|---|---|---|---|
| CD3 | AF532 | UCHT1 | LifeTech | 58-0038-42 |
| CD4 | BV605 | RPA-T4 | BD Biosciences | 562658 |
| CD8 | BUV496 | RPA-T8 | BD Biosciences | 612942 |
| CD14 | V500 | M5E2 | BD Biosciences | 561391 |
| CD19 | V500 | HIB19 | BD Biosciences | 561121 |
| CD137 | APC | 4B4-1 | Biolegend | 309810 |
| CD134/OX40 | PE-Cy7 | Ber-ACT35 | Biolegend | 350012 |
| CD69 | PE | FN50 | BD Biosciences | 555531 |
| CD45RA | BV421 | HI100 | Biolegend | 304130 |
| CCR7 | FITC | G043H7 | Biolegend | 353216 |
| Live/Dead Viability | eF506/Aqua | - | Invitrogen | 65-0866-18 |
